# Mitosis Detection in the Wild Using Detection Transformers

**DOI:** 10.1101/2025.09.03.673198

**Authors:** Vidushi Walia, Dhandapani Nandagopal, Sujatha Kotte, Vangala Govindakrishnan Saipradeep, Thomas Joseph, Naveen Sivadasan, Bhagat Singh Lali

## Abstract

**Background:** Identification of mitotic cells and its down-stream analysis, is an important parameter in understanding the pathology of cancer, predicting response to chemotherapy and overall survival. However, their reliable detection remains challenging due to morphological overlap with other cellular structures, resulting in variability and high levels of false positives. Current artificial intelligence (AI) algorithms further face limitations when confronted with tissue heterogeneity and often underperform in non-tumor, inflamed, or necrotic regions. To address these challenges, Track 1 of MIDOG 2025 challenge expands the scope of mitotic figure detection to include all tissue regions, encompassing both hotspot and non-hotspot areas, thereby promoting real-world clinical applicability.

**Methods:** We propose a generalized deep learning based mitotic detection model (MD model) for robust and accurate detection of mitotic figures using the MIDOG-2025 challenge dataset. Our model enables robust and accurate detection of mitotic figures and demonstrates strong generalization to domain shifts arising from diverse histological regions, variations in scanners, tumor subtypes and laboratory protocols.

**Results:** Our approach showed a consistent performance with an F1-score of 0.7769 with a high recall of 0.8222. Our approach outperforms the baselines and generalizes well across the tumor types on the preliminary test set.

**Conclusion:** Our approach makes the final predictions with reduced false positives and improved detection accuracy. It generalizes well to address the domain shifts caused by diverse histological regions across the different tumor types among others.

## Introduction

The conventional approach for identifying mitotic cells relies on manual examination of Hematoxylin and Eosin (H&E)-stained tissue slides. Recent advances in digital pathology(1) have enabled the use of high-resolution scanners to capture images from whole slide images (WSIs). Despite these developments, state-of-the-art deep learning (DL) models trained on WSIs are often restricted to specific tumor tissue regions, particularly mitotic hotspots. As a result, their performance degrades significantly when applied to non-hotspot or non-tumor regions, where morphological variability is higher and mitotic figures are sparse. This domain shift between hotspot and non-hotspot regions introduces considerable challenges, leading to reduced generalizability of existing models. This challenge is further compounded by inter-tumor variability, as mitotic morphology can differ substantially across cancer types, thereby limiting model generalization. Additionally, inter-laboratory differences in staining protocols and scanner characteristics exacerbate these issues. Together, these factors hinder the reliability and clinical adoption of computer-assisted mitosis detection methods.(2)

To help address the different shortcomings related to automated detection of mitotic figures, there have been multiple efforts in the past through the form of Challenges like TUPAC16 (3), Mitos & Atypia 14 (4), MIDOG-2021 (5) and MIDOG-2022 (6). Track 1 of the MIDOG-2025 Challenge (7) looks further to develop algorithms for the accurate detection of mitotic figures across a wide range of tissue regions. These regions include the tumor hotspot areas as well as the more challenging regions such as the non-tumor areas, inflamed tissue, staining artifacts and necrotic regions. In this work, we propose a deep learning-based model that detects mitotic figures with improved generalization performance.

## Material and Methods

Fig. 1 and Fig. 2 architecture of our two proposed strategies for Mitosis detection, namely, Ensemble Model and MD Model respectively.

**Fig. 1.**
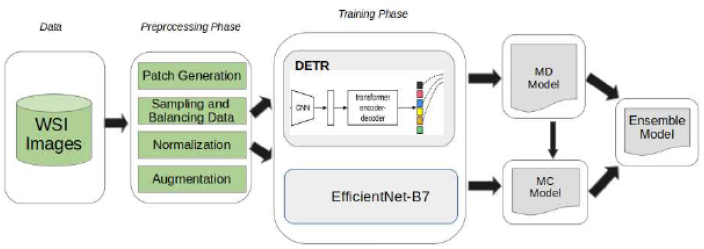
Architecture of Ensemble Model

**Fig. 2.**
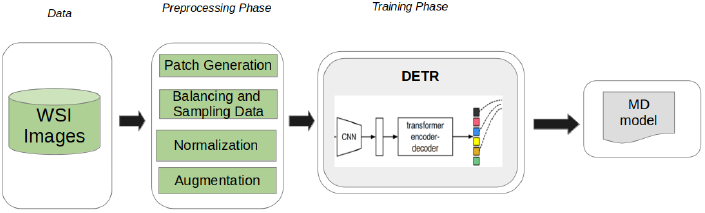
Architecture of MD Model

### Data

Our model was trained and evaluated using three mitotic figure (MF) detection datasets that span a broad range of domains. The primary dataset, MIDOG++ (8) , consists of region-of-interest images derived from 503 histopathology specimens across seven tumor types, including human breast cancer, neuroendocrine tumor, melanoma, canine mast cell tumor, lung cancer and lymphoma, providing 11,937 annotated MFs. These samples originate from multiple laboratories and scanners, thereby introducing substantial domain variability. In addition, we utilized two additional canine tumor datasets: the Canine Cutaneous Mast Cell Tumor (CCMCT) (9) dataset , comprising 32 whole-slide images of mast cell tumors with exhaustive mitotic annotations, and the Canine Mammary Carcinoma (CMC)(10) dataset , which includes 21 whole-slide images of mammary carcinoma with detailed MF annotations. Both datasets introduce species- and tissue-type domain shifts, complementing the human data.

### Data Pre-processing and Augmentation

Train and validation datasets were created using 80:20 split on the 556 annotated images. Based on the labels of the training data, 512x512 pixel contextual regions around the mitotic/imposter figures were extracted from each WSI. Multiple contextual regions were extracted for each labeled figure after applying random shift to the context window. Standard augmentation methods such as horizontal flipping, vertical flipping, random rescaling, random cropping, and random rotation are performed to make the model invariant to geometric perturbations. Moreover, Random HSV is also adopted to randomly change the hue, saturation, and value of images in the hue saturation value (HSV) color space, making the model robust to color perturbations.

### DETR based object detection

Detection Transformer (DETR) (11) was trained on the MIDOG 2025 dataset. Based on the F1 score on the validation dataset, Resnet50-DC5 backbone pretrained on ImageNet was selected from among the DETR backbones namely, Resnet101, Resnet50 and Resnet101-DC5. The hyperparameters of DETR were chosen using Optuna hyperparameter optimization framework (12). For the test WSI images, using a sliding window with overlay of 50 pixels, cropped regions of size 512x512 pixels were extracted. Each cropped region was fed to the trained DETR model. The DETR output consisted of the identified objects and their associated class probabilities. All mitotic predictions with class probability greater than or equal to 0.85 were directly included in the output with mitotic class label. For the preliminary test phase as well as the final test phase, we trained the DETR model on the entire labelled training data of 556 images. We trained our DETR-based object detection framework using two distinct strategies. The first is a straightforward detection model built on DETR-architecture, which we refer to as the MD Model. The second strategy adopts an ensemble approach, where we combine the MD Model with an independently trained classifier to produce the final Ensemble Model.

### DAB-DETR based object detection

We trained DAB-DETR (13) on the challenge dataset and explored two strategies. In the first, we directly employed the DAB-DETR architecture for mitotic figure detection. We name this strategy as Vanilla DAB-DETR. In the second, termed as Frozen DAB-DETR, we integrated embeddings from a foundation model to guide the training of DAB-DETR transformer.

### Ablation Studies

We conducted extensive ablation studies to understand the effect of different parameters on the final model performance. This included experiments on different patch sizes (512 vs 128 pixels), stain augmentation vs stain normalization, choice of DETR backbones (ResNet50, ResNet50-DC5, ResNet101 and ResNet101-DC5), Classifier CNN models (ResNet, Densenet, EfficientNet-B7, EfficientNet-B8. EfficientNet-B7) and deep supervision where Hematoxylin channel was added as a fourth channel to the images in addition to RGB channels. On the basis of these experiments, we chose a patch size of 512 pixels, Random HSV for stain augmentation, ResNet50-DC5 as DETR CNN backbone and EfficientNet-B7 as the classifier. However, adding a separate Hematoxylin channel did not show any improvement in the performance on the validation set. All experiments are repeated under the same training and evaluation conditions.

## Results

### Metrics

Submissions to the MIDOG 2025 Challenge were evaluated across multiple performance metrics, including precision, recall, F1 score, average precision, and FROC/AUC. In addition, domain specific F1 score were reported for each of the four tumor types comprising the preliminary test set.

Table 2 shows the performance of our different approaches in comparison with the challenge baseline across the evaluated metrics. Two of our approaches, MD Model and Ensemble Model, demonstrated promising results, surpassing the baseline in most of the metrics. The MD Model and Ensemble Model obtained F1-scores of 0.7769 and 0.7819 respectively, surpassing the baseline F1 score of 0.7672, reflecting competitive performance on the leaderboard. This pattern of performance was observed across the metrics, where MD Model and Ensemble model demonstrated either comparable or superior performance across the remaining evaluation metrics, indicating robustness of our approach. Detailed results on the performance of our submitted approaches can be found in Table 2.

**Table 1.** Shows the number of patches and annotation of Mitotic figure from the three dataset used for training.

|  | MIDOG++ |  | CMC |  | CCMCT |  |
| --- | --- | --- | --- | --- | --- | --- |
|  | # Patches | # Mitotic Figures | # Patches | # Mitotic Figures | # Patches | # Mitotic Figures |
| **Train/Eval** | 62219 | 55615 | 51406 | 30438 | 33988 | 41669 |

**Table 2.** Depicts the performance of our various approaches and the baseline provided by the challenge on preliminary test dataset for Track 1 of MIDOG 2025 challenge.

|  | Precision | Recall | F1- score | Avg. Precision | FROC/AUC | F1 Domain 1 | F1 Domain 2 | F1 Domain 3 | F1 Domain 4 |
| --- | --- | --- | --- | --- | --- | --- | --- | --- | --- |
| **MD Model** | 0.7363 | **0.8222** | 0.7769 | **0.8118** | **5.8088** | 0.7765 | 0.7837 | **0.7626** | **0.8000** |
| **Ensemble Model** | **0.7724** | 0.7917 | **0.7819** | 0.8108 | 5.7974 | 0.7848 | 0.7727 | 0.7895 | 0.7895 |
| **Vanilla DAB-DETR** | 0.7184 | 0.5527 | 0.6248 | 0.4959 | 4.0653 | 0.6585 | 0.5514 | 0.6944 | 0.6250 |
| **Frozen DAB-DETR** | 0.7101 | 0.6805 | 0.6950 | 0.6274 | 4.639 | 0.7857 | 0.6083 | 0.7157 | 0.8219 |
| **Challenge Baseline** | 0.7441 | 0.7917 | 0.7672 | 0.8001 | 5.6613 | **0.7952** | **0.8115** | 0.7055 | 0.7778 |

Table 2 also shows performance of various approaches and baseline of the different tumor types. F1 score of different Domains demonstrate that our approach showed comparable or superior performance across four different tumor types. The MD Model and Ensemble Model both showed comparative performance. Among them, we selected MD Model due to its computational efficiency. Considering the timing constraints of the submission, MD Model turned out to be our ultimate choice for final submission.

Figure 3 presents a selection of image patches with overlaid bounding boxes for both the ground truth and the predictions from the MD Model. The left side visual fields represent the overlaid ground truth while the right side visual fields show the overlaid predictions from MD Model.

**Fig. 3.**
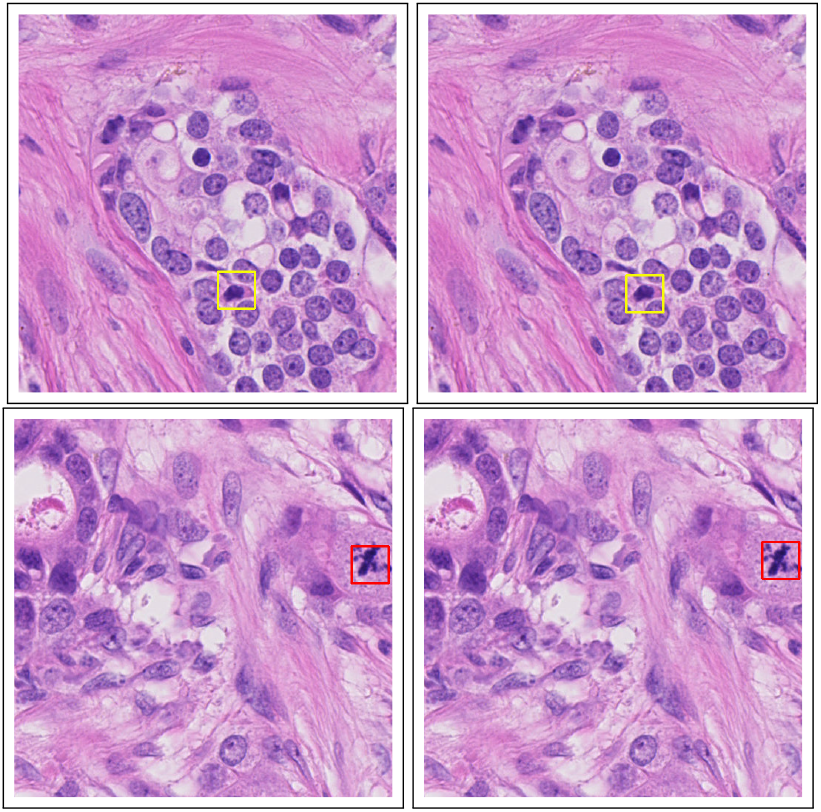
An example of visual fields overlaid with Mitotic Figure (red) and Non Mitotic Figure (yellow). Left hand side images show the overlay of the ground truth while right hand side images show the overlay of predictions from our MD model.

## Discussion

Our work demonstrates that Vision Transformer-based detection offers a robust solution to mitosis identification in histology images. Our proposed framework generalizes effectively across variations in tissue regions (hotspot vs non-hotspot regions), staining, scanners, laboratory protocols and tumor types.

DAB-DETR and its frozen variant represent a promising direction for our future work. We plan to further refine these approaches to enhance their capability for mitotic figure detection, with a particular focus on improving generalization across diverse domains. Incorporating stronger domain adaptation strategies and optimizing the integration of foundation model embeddings will be our central focus.

## ACKNOWLEDGEMENTS

The authors thank the organizers of MIDOG-2025 for sharing the data with the community.

## Supplemental Materials: Mitosis Detection in the Wild Using Detection Transformers

Additional results on the ablation studies on various augmentation startegies are shown in the table below. For this comparison we use our internal validation data.

**Table S3.**
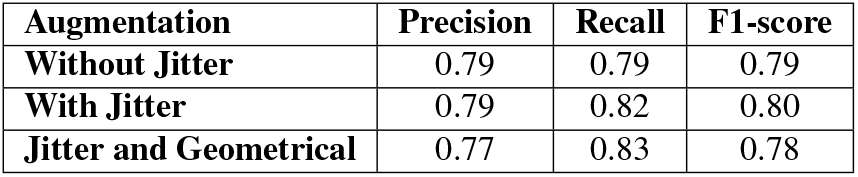
Depicts the performance of our various albumentation approaches as part of ablation studies on our internal validation set.

| Augmentation | Precision | Recall | F1-score |
| --- | --- | --- | --- |
| Without Jitter | 0.79 | 0.79 | 0.79 |
| With Jitter | 0.79 | 0.82 | 0.80 |
| Jitter and Geometrical | 0.77 | 0.83 | 0.78 |

## Bibliography

1. H. Dawson. Digital pathology–rising to the challenge. Frontiers in medicine, 9:888896, 2022.

2. G. Jiménez and D. Racoceanu. Deep learning for semantic segmentation vs. classification in computational pathology: application to mitosis analysis in breast cancer grading. Frontiers in bioengineering and biotechnology, 7:145, 2019.

3. M. Veta, Y.J Heng, N. Stathonikos, B.E Bejnordi, F. Beca, T. Wollmann, K. Rohr, MA. Shah, D. Wang, M. Rousson, M. Hedlund, D. Tellez, F. Ciompi, E. Zerhouni, D. Lanyi, M. Viana, V. Kovalev, V. Liauchuk, H.A Phoulady, T. Qaiser, S. Graham, N. Rajpoot, E. Sjöblom, J. Molin, K. Paeng, S. Hwang, S. Park, Z. Jia, E.I Chang, Y. Xu, A.H Beck, P.J van Diest, and J.P.W Pluim. Predicting breast tumor proliferation from whole-slide images: The tupac16 challenge. Medical Image Analysis, 54:111–21, 2019.

4. Mitos-atypia-14 - grand challenge. https://mitos-atypia-14.grand-challenge.org/, 2014.

5. M. Aubreville, N. Stathonikos, C.A Bertram, R. Klopfleisch, N.. Ter Hoeve, F. Ciompi, F. Wilm, C. Marzahl, T.A Donovan, A. Maier, J. Breen, N. Ravikumar, Y. Chung, J. Park, R. Nateghi, F. Pourakpour, R.H.J Fick, S. Ben Hadj, M. Jahanifar, A. Shephard, J. Dexl, T. Wittenberg, S. Kondo, M.W Lafarge, V.H Koelzer, J. Liang, Y. Wang, X. Long, J. Liu, S. Razavi, A. Khademi, S. Yang, X. Wang, R. Erber, A. Klang, K. Lipnik, P. Bolfa, M.J Dark, G. Wasinger, M. Veta, and K. Breininger. Mitosis domain generalization in histopathology images - the midog challenge. Medical Image Analysis, 84:102699, 2023.

6. M. Aubreville, C. Bertram, K. Breininger, S. Jabari, N. Stathonikos, and M. Veta. Mitosis domain generalization challenge 2022. In 25th International Conference on Medical Image Computing and Computer Assisted Intervention (MICCAI 2022), 2022. doi: 10.5281/zenodo.6362337.

7. J. Ammeling, M. Aubreville, S. Banerjee, C.A Bertram, K. Breininger, D. Hirling, P. Horvath, N. Stathonikos, and M. Veta. Mitosis domain generalization challenge 2025. 10.5281/zenodo.15077361, 2025.

8. M. Aubreville, F. Wilm, N. Stathonikos, K. Breininger, T.A Donovan, S. Jabari, M. Veta, J. Ganz, J. Ammeling, P.J van Diest, et al. A comprehensive multi-domain dataset for mitotic figure detection. Scientific data, 10(1):484, 2023.

9. C.A Bertram, M. Aubreville, C. Marzahl, A. Maier, and R. Klopfleisch. A large-scale dataset for mitotic figure assessment on whole slide images of canine cutaneous mast cell tumor. Scientific data, 6(1):274, 2019.

10. M. Aubreville, C.A Bertram, T.A Donovan, C. Marzahl, A. Maier, and R. Klopfleisch. A completely annotated whole slide image dataset of canine breast cancer to aid human breast cancer research. Scientific data, 7(1):417, 2020.

11. N. Carion, F. Massa, G. Synnaeve, N. Usunier, A. Kirillov, and Zagoruyko S. End-to-end object detection with transformers. ECCV, 2020.

12. T. Akiba, S. Sano, T. Yanase, T. Ohta, and M. Koyama. Optuna: A next-generation hyperparameter optimization framework. KDD ‘19: Proceedings of the 25th ACM SIGKDD International Conference on Knowledge Discovery Data Mining, pages 2623 – 2631, 2019.

13. Shilong Liu, Feng Li, Hao Zhang, Xiao Yang, Xianbiao Qi, Hang Su, Jun Zhu, and Lei Zhang. Dab-detr: Dynamic anchor boxes are better queries for dextr. arXiv preprint 2201.12329, 2022.

